# RepairSig: Deconvolution of DNA damage and repair contributions to the mutational landscape of cancer

**DOI:** 10.1101/2020.11.21.392878

**Authors:** Damian Wojtowicz, Jan Hoinka, Bayarbaatar Amgalan, Yoo-Ah Kim, Teresa M. Przytycka

**Author notes:** Corresponding authors. (D.W.), (T.M.P.). First authors (Damian Wojtowicz, Jan Hoinka).

## Abstract

Many mutagenic processes leave characteristic imprints on cancer genomes known as mutational signatures. These signatures have been of recent interest regarding their applicability in studying processes shaping the mutational landscape of cancer. In particular, pinpointing the presence of altered DNA repair pathways can have important therapeutic implications. However, mutational signatures of DNA repair deficiencies are often hard to infer. This challenge emerges as a result of deficient DNA repair processes acting by modifying the outcome of other mutagens. Thus, they exhibit non-additive effects that are not depicted by the current paradigm for modeling mutational processes as independent signatures. To close this gap, we present RepairSig, a method that accounts for interactions between DNA damage and repair and is able to uncover unbiased signatures of deficient DNA repair processes. In particular, RepairSig was able to replace three MMR deficiency signatures previously proposed to be active in breast cancer, with just one signature strikingly similar to the experimentally derived signature. As the first method to model interactions between mutagenic processes, RepairSig is an important step towards biologically more realistic modeling of mutational processes in cancer. The source code for RepairSig is publicly available at https://github.com/ncbi/RepairSig.

## 1 Introduction

Studies of cancer mutations have typically focused on identifying cancer-driver mutations. However, cancer genomes also accumulate a large number of somatic mutations resulting from various endogenous and exogenous causes, including spontaneous deamination of cytosines, mutations triggered by carcinogenic exposures, or cancer-related aberrations of the DNA maintenance machinery. These mutations are typically harmless passenger mutations, but analysis of their patterns can provide useful information regarding mutational processes acting on cancer genomes. Indeed, different mutagenic processes often leave characteristic mutation imprints in cancer genomes [6]. Identifying processes shaping the mutational landscape of cancer is an important step towards understanding tumorgenesis and has the potential to inform therapeutic and preventative measures. Consequently, the development of methods intended for the identification of mutation patterns present in cancer genomes and linking such patterns to specific mutagenic processes has become a research topic of broad interest [7].

Approaches for *de novo* discovery of mutational signatures typically aim to define an optimal set of signatures so that mutation counts observed in all samples can be expressed as the sum of contributions of individual signatures weighted by the patient-specific signature activity (i.e. exposure). These methods include the pioneering work of Alexandrov et al. utilizing non-negative matrix factorization (NMF) [6,7], as well as subsequent approaches that incorporate various techniques to promote sparsity or impose other constraints on the model [9,15,19,25,27]. The fundamental objective in delineating mutational signatures consists in leveraging this information to study the mutational processes contributing to the mutations observed in individual cancer genomes. Indeed, many so-called COSMIC mutational signatures, derived by Alexandrov et al. [5,6] and used as the reference signatures in many other studies, have been successfully assigned to specific mutational processes. Elucidating mutagenic causes of signatures with unknown etiology is a subject of intense computational and experimental studies [8,16,24,28,31].

Concurrently with these advances, our understanding of the interplay between mutagenic processes and their signatures is increasingly shifting away from a one-to-one correspondence model, demanding the development of models incorporating more complex relationships. This change is mainly driven by mounting evidence suggesting that the widely used assumption of additivity of mutagenic processes is an oversimplification [12,28,29]. For example, deficiency in mismatch repair (MMR), a DNA repair process recognizing and correcting mismatched (non-complementary) nucleotides in the otherwise complementary paired DNA strands, has been indicated in at least eight different COSMIC signatures. Yet, our recent analysis, utilizing the so-called signature RePrint [29], suggested that these signatures are likely to represent composite mutagenic processes where MMR dysfunction is superimposed on different types of “initial” DNA damage. As a case in point, a recent study convincingly demonstrated that two of these signatures are indeed composite signatures in which different types of DNA damage (caused by mutations in polymerases POLE and POLD1) are accompanied by a common deficiency in mismatch repair mechanism [13].

Despite the growing evidence indicating that many current signatures associated with DNA damage represent a combination of factors related to DNA damage and repair, separating the contribution of DNA repair deficiency from the primary DNA damage has remained an understudied research topic. While RePrint is capable of identifying signatures that are likely to share common DNA repair deficiency mechanisms, it cannot provide such a decomposition. Therefore, the contributions from deficiencies of DNA repair processes to other mutational signatures are yet to be fully explored. Understating altered DNA repair pathways can have important therapeutic implications for personalized treatments. A relevant instance concerns patients with homologous recombination deficiency (HRD) responding to drugs known as PARP inhibitors. Moreover, leveraging other relations between DNA repair deficiency and drug action may also improve the efficacy of cancer treatments [21].

To close this gap, we propose RepairSig, a new computational approach that accounts for the non-additivity of DNA damage and repair processes. More precisely, RepairSig explicitly models the composition of (i) *primary mutagenic processes*, that correspond to DNA damage processes with normally functioning DNA repair mechanism (Fig. 1A, left) and (ii) *secondary mutagenic processes*, which correspond to the deficiency of the DNA repair mechanism, i.e. processes altering the way in which the errors introduced by the primary processes are repaired (Fig. 1A, right). We assume that the signatures of the primary processes are known in the model while signatures of the secondary processes are to be inferred. In addition, our framework design is flexible enough to allow for the inference of primary mutational signatures as well.

**Fig. 1.**
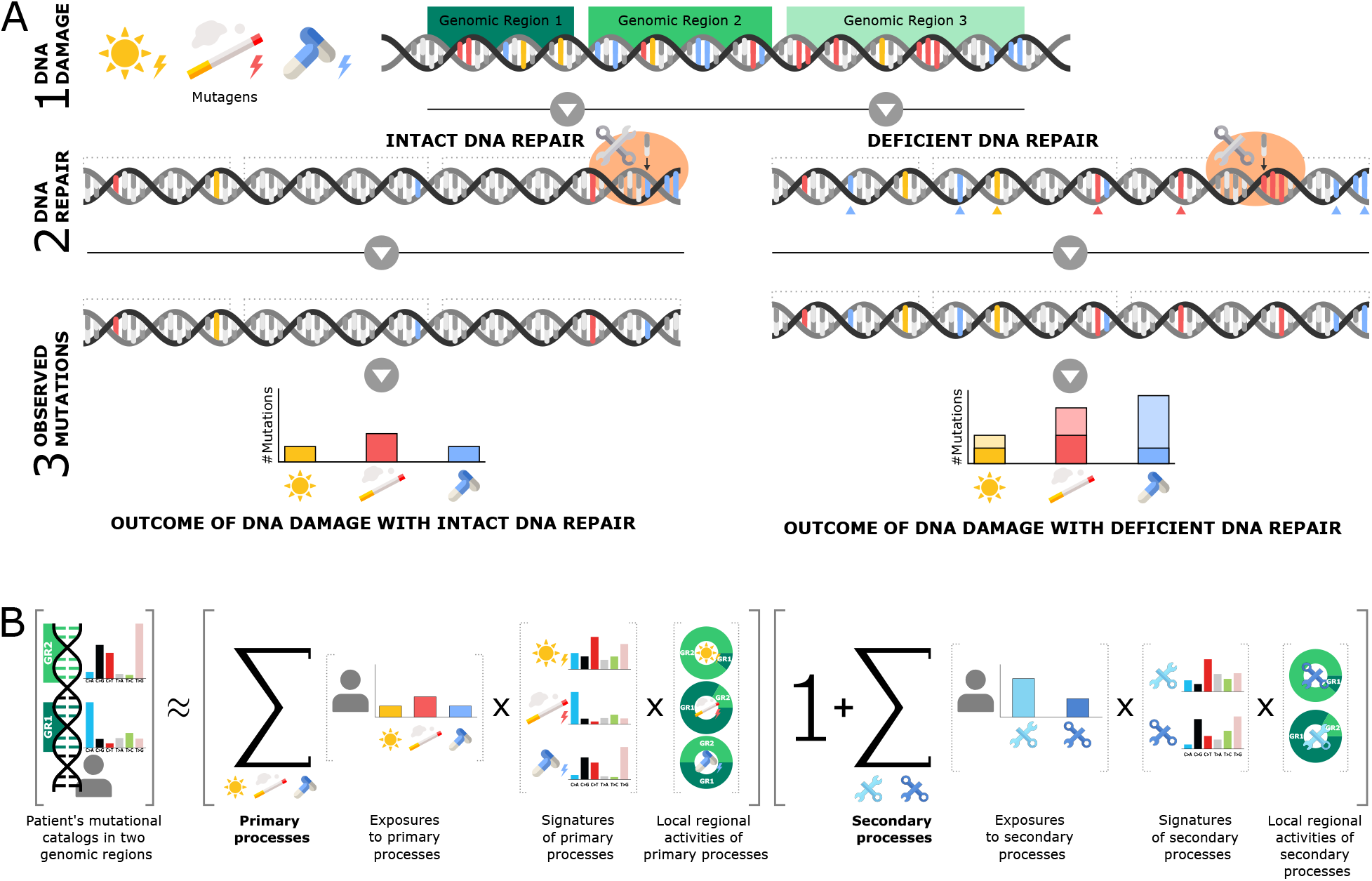
Method overview. (**A**) **Interplay between DNA damage processes and DNA repair mechanisms.** DNA is exposed to exogenous and endogenous mutagenic processes that cause DNA damage. The amount of DNA primary damage (lesions) can vary between genomic regions and it depends on the level of exposure to DNA damage source, the DNA sequence context and state. DNA repair mechanism serves to counteract the primary damage. However, even the intact DNA repair system is not a perfect mechanism and some primary damage is improperly repaired or entirely missed by the repair mechanism leading to this damage being observed as mutations (left). Furthermore, in some individuals, the DNA repair mechanism may become defective, for example, due to a mutation in a DNA repair gene. Consequently, DNA damage tends to accumulate resulting in mutations that would normally have been repaired under a fully functional repair system (right). (**B**) **Overview of RepairSig**. Its architecture is illustrated for one patient’s genome, two genomic regions (indicated in green), three primary processes (UV light, smoking, and drug toxicity), and two secondary processes (blue repair symbols). The cancer genome can be represented as an interplay between the primary and secondary processes, where the latter amplify DNA damage caused by the former. Mutations attributed to each type of these processes can be expressed in terms of their exposures, mutational signatures, and local regional activities. The mutations of all types caused by the primary mutational processes are modeled as a linear superposition of the respective exposures, signatures and local activities of the mutagenic processes (see first summation term). The amplification of mutations due to the activity of the secondary mutational processes is expressed in a similar way (second summation term). Finally, the total mutational catalog of the cancer genome consists of the primary processes’ mutations whose number is further augmented by the secondary processes concerning deficient DNA repair mechanism. Please note that “ **x**” does not represent a regular matrix multiplication, see Eq. (2) for the formal problem formulation.

The aforementioned interactions between DNA damage and repair processes are not the only non-additive processes that might influence the mutation landscape. Besides the interplay between DNA damage and repair, the distribution of mutations across the genome is additionally influenced by the so-called *mutation opportunity* – the distribution of sites that are vulnerable to damage or predisposed to repair. Most existing models operate under the assumption that even though mutation opportunities might vary within the genome, these variations are identical in different cancer genomes. However, mutation opportunities can arise differently in distinct genomes in response to DNA stress, double-strand breaks, shifts in phosphorylation patterns, or other endogenous cancer-related processes. To account for these aspects, we designed our model to allow for the detection of shifts in mutation opportunity due to large-scale features such as chromatin organization, transcription orientation, replication timing and direction, as well as local chromatin features including transcription factors, nucleosomes, and non-B DNA motifs [11].

Mathematically, the model behind RepairSig can be conveniently defined in terms of the exposures, signatures, and local activities of the primary and secondary processes (see Fig. 1B and Eq. (2)). However, finding a satisfactory solution for this problem formulation is not straightforward. Thus, for efficient model optimization, we translate the formulation into higher-order tensor algebra (see Eq. (4)) and leverage the powerful TensorFlow optimization API [4] to find solutions for a given input.

We first tested our approach using simulated data and show that RepairSig is capable of recovering the correct decomposition in a variety of scenarios. Subsequently, we applied our model to cancer data and found that the MMR deficiency signature obtained using RepairSig is in better agreement with the experimental signature obtained with CRISPR-Cas9 based knockout compared to any other COSMIC signature related to MMR. Our method also allowed us to obtain interesting insights into mutagenesis in breast cancer. To the best of our knowledge, RepairSig is the first approach to infer DNA repair deficiency signatures that builds on a biologically realistic, non-additive, model-based composition of mutagenic processes.

## 2 Results

Here, we present a novel method, RepairSig, to model DNA damage and repair processes in cancer. Our approach strives to explain the catalogs of somatic mutations observed in cancer genomes as a joint product of DNA damage and repair processes while accounting for additional factors that underlie the activities of these processes. As a proof-of-principle, we applied RepairSig to breast cancer genomes and other cancers known to be affected by the impaired DNA repair mechanism.

### 2.1 A combined model for primary and secondary mutagenic processes

#### Preliminaries

We assume that somatic mutations in cancer fall into *K* categories. Typically, *K* = 96, corresponding to 6 types of mutation base substitutions and 16 combinations of flanking nucleotide residues (e.g. TCC>TTC represents C>T mutation in T_C nucleotide context). These mutations are presumed to be the result of the activity of *N* primary mutational processes and *J* secondary mutational processes. *Primary mutagenic processes* typically correspond to classical exogenous processes such as tobacco smoking, UV light exposure, and other processes that are considered to be independent of each other (Fig. 1A, left). Endogenous processes that introduce DNA damage in an independent manner are also included in this group. On the other hand, *secondary mutational processes* modify the outcome of primary processes (Fig. 1A, right). One important example of such a process is the DNA mismatch repair (MMR) deficiency discussed in the Introduction. The deficiency of the MMR mechanism does not cause new mutations by itself but leads to accelerated accumulation of improperly repaired damage caused by primary processes. The activity of primary and secondary processes can be influenced by additional genomic factors such as transcription and replication status, chromatin structure, or DNA shape. The number of different domains associated with these factors is denoted by *L*. The goal of RepairSig is to explain catalogs of somatic mutations of *K* types observed in *G* cancer genomes as the joint product of *N* primary and *J* secondary processes accounting for *L* genomic domains that underlie mutation rates of these processes (Fig. 1B).

#### Model specification

Formally, let *M* be a mutation count matrix computed from sequenced cancer genomes, i.e. a three-dimensional matrix of size *G* × *K* × *L* containing the mutational profiles from each of the *L* genomic regions in *G* cancer genomes. In turn, each profile contains the mutation frequencies of *K* mutation types computed from the respective patient’s catalog of mutations for each type of genomic region. We model mutational signatures with two signature matrices *P* (*N* × *K*) and *Q* (*J* × *K*) for *N* predefined primary and *J* unknown secondary mutational signatures, respectively. These matrices specify the probabilities of generating each of the *K* mutation types by each mutational process. The overall activities of the primary and secondary mutational processes are represented by two non-negative matrices *A* (*G* × *N*) and *D* (*J* × *N*). They contain, for each of the *G* genomes, the patient’s genome-wide exposure to each of *N* primary and *J* secondary processes, respectively. Finally, the local regional activity (mutation rates) of these processes across *L* predefined genomic regions are represented by non-negative matrices *W* (*N* × *L*) and *R* (*J* × *L*). Here, each row corresponds to relative mutation rates with which a mutational process operates in these regions, and each row models a probability distribution over *L* types of genomic regions.

If all mutational processes are primary (Fig. 1A, left), *m_g,k,l_*, i.e. the number of mutations of the *k*-th mutation type in the *l*-th region of genome *g*, can be expressed as 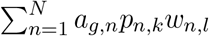, the sum over these processes weighted by their exposures and local region-specific activity strength. However, if the DNA repair process is defective, the number of mutations can be amplified by the secondary mutational processes leading to excessive accumulation of mutations with respect to the intact repair mechanism (Fig. 1A, right). In particular, a signature of a secondary process does not describe the frequency of introducing new mutations but rather the frequency of not repairing a lesion that normally would have been repaired if the process was fully functional. Consistently, in RepairSig, the primary and secondary processes are not treated as independent contributions but the secondary processes act on the outcome of the primary processes. Such an additional acquisition of mutations caused by the *j*-th secondary process can be expressed by a multiplicative factor of *d_g,j_q_j,k_r_j,l_*. Finally, the joint effect of all *N* primary and *J* secondary mutational processes, their exposure, and local regional strengths on the observed number of mutations of the *k*-th mutation type in the *l*-th region of genome *g* can be expressed as shown in Eq. (1) and further approximated by Eq. (2) after removing second-order terms.

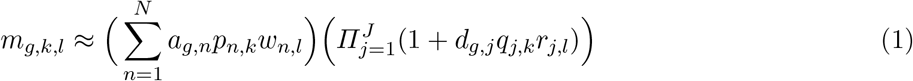

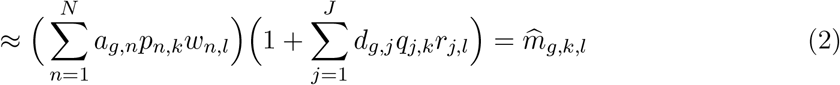

RepairSig seeks to identify non-negative matrices *W*, *A*, *Q*, *R*, and *D* that minimize the difference (Frobenius norm) between the mutation count matrix *M* and its approximation 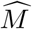 following Eq. (2). Formally, we solve the following optimization problem:

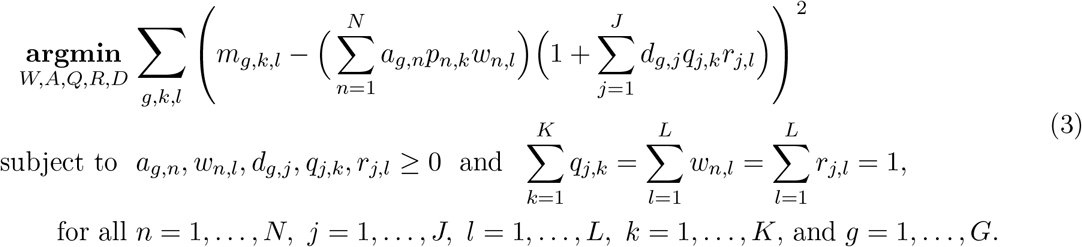

Additional details of the model, its formulation using tensor algebra, implementation details, and optimization strategy are described in Materials and Methods.

#### Model evaluation

We validated our method on simulated and real cancer data, demonstrating that it allows for accurate inference of secondary signatures as well as signature exposures and their local activity strengths. First, we tested RepairSig on the simulated data for a varying number of the secondary mutational signatures (see Supplementary Materials). Supplementary Fig. S1 highlights RepairSig’s ability to recover all matrices *W*, *A*, *Q*, *R*, *D*, and *M* with high accuracy – the average Pearson’s correlation between the simulated and inferred values of each matrix was higher than 0.97, 0.96, and 0.86 for *J* = 1, 2, 3, respectively.

We also evaluated RepairSig on BRCA whole-genome cancer sequences with local regional activity modeled using two types of genomic data, namely transcriptional strands and replicative time domains (see Materials and Methods). To select the appropriate number of secondary signatures in RepairSig, we perform 100 optimizations for increasing numbers of signatures (*J* = 1,…, 4) while initializing the matrices at random for each optimization. The best performance was achieved for *J* = 2 by balancing the reconstruction error and the signature reproducibility (Supplementary Fig. S2A). Moreover, RepairSig (*J* = 2) outperforms additive models in terms of the reconstruction error, including those with more free parameters than our model (Supplementary Fig. S2B). Note that these additive models assume all mutational processes to be primary and therefore cannot account for any interactions between DNA damage and repair processes in the cancer data. Similar results were obtained for other cancer types studied here (data not shown). Thus, we adopted the RepairSig model with *J* = 2 for the analyses presented in this study.

### 2.2 DNA mismatch repair deficiency in breast cancer can be explained by a single MMRd signature

First, we applied RepairSig to whole-genome breast cancer (BRCA) data (see Materials and Methods). Previous studies identified the activity of at least 12 COSMIC signatures in this cancer including at least 3 signatures associated with MMR deficiency (MMRd) [23,14,5]. We hypothesized that this large number of MMRd related signatures is a result of interactions between MMR deficiency process and other mutational processes active in BRCA. Thus, RepairSig should require a much smaller number of MMRd signatures as it models interactions between DNA damage and repair processes. As the primary signatures in RepairSig, we used COSMIC signatures previously shown to be present in BRCA [23], excluding the 3 signatures associated with MMRd (Signatures 6, 20, and 26). In addition, we chose two distinct partitions of the genome into local genomic regions defined by the strand-specific direction of gene transcription and discretized replication timing data (see Materials and Methods). In each setting, RepairSig inferred two secondary signatures (Fig. 2A,B). Interestingly, both models produced one mutual signature (RepairSig.T1 and RepairSig.R1). This common signature resembles the *Δ*MSH6 signature derived from the CRISPR-Cas9-based knockout of the MMR gene MSH6 (cosine similarity > 0.9) [31]. The remaining signatures did not show any mutual similarity and, as we argue below, are most likely not directly related to MMR deficiency but rather a different secondary process. Thus, as hypothesized, RepairSig was able to identify a single MMRd related secondary signature replacing three COSMIC MMRd signatures present in BRCA. The classical additive model requires more free parameters to achieve similar performance to RepairSig (Supplementary Fig.S2B). In addition, the identified signature corresponds very well to the experimentally derived MMRd signature of *Δ*MSH6.

**Fig. 2.**
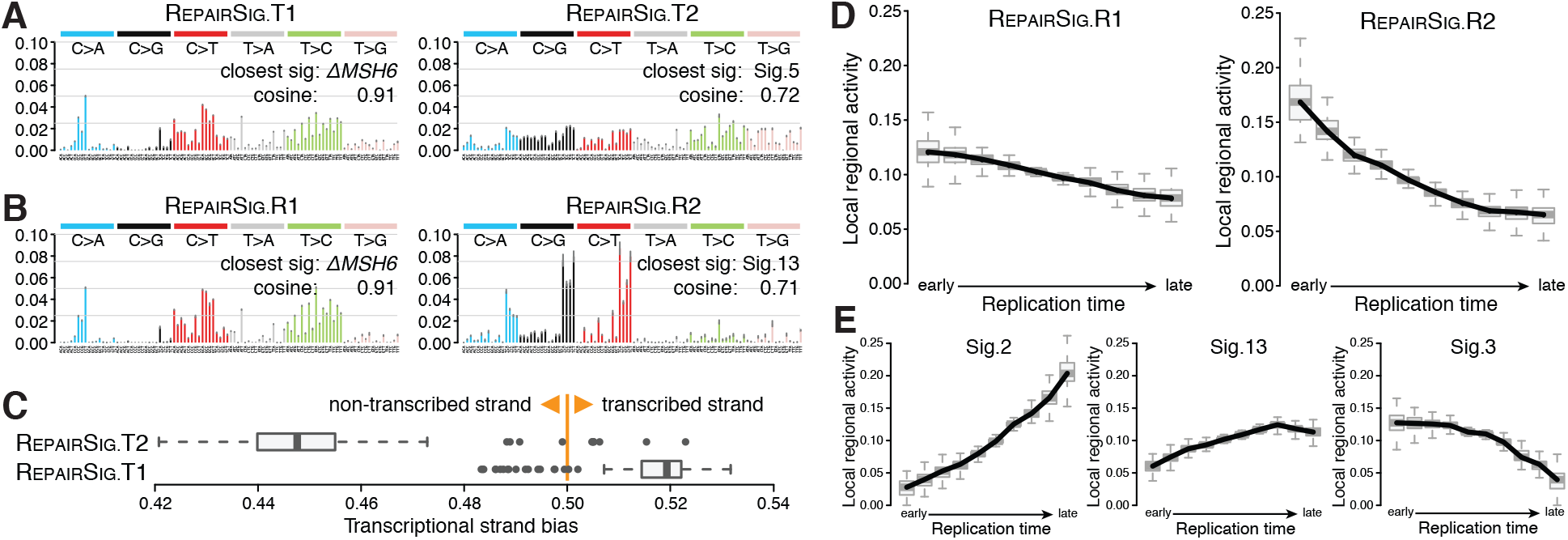
Results of RepairSig in BRCA. (**A,B**) The mutational signatures inferred by RepairSig from cancer sequencing data divided into genomic regions based on transcriptional strand orientation (**A**) and replication timing domains (**B**). The closest known signature and its cosine similarity are shown for each RepairSig signature. (**C**) Transcriptional strand bias, i.e. an asymmetry in the number of mutations found on transcribed and non-transcribed strands, for the transcription-based RepairSig signatures. (**D,E**) Local regional activity distribution of the replication-based RepairSig signatures (**D**) and Signatures 2, 13, and 3 (**E**) over replication time domains modeled by RepairSig.

The remaining two secondary signatures RepairSig.T2 and RepairSig.R2 also provide important insights. RepairSig.T2 is relatively flat and is most similar to COSMIC Signature 5 (Fig. 2A) that is associated with aging. This signature also emerges in cancers with a deficiency in the nucleotide excision repair (NER), another key DNA repair mechanism. Unfortunately, no experimentally inferred ground truth signature for NER deficiency has been established to date. However, RepairSig.T2 additionally resembles the “flat” signature obtained from the CRISPR-Cas9-based knockout of the FANCC gene involved in several DNA repair pathways including NER [22,26] (cosine similarity: 0.7). Since this similarity is weaker compared to the MMRd signature, we leveraged the local regional activities inferred by RepairSig for subsequent analysis to gain additional insight. A striking property of RepairSig.T2 is its exceptionally high transcriptional strand bias, i.e. an asymmetry in the number of mutations found on the transcribed and non-transcribed strand (Fig. 2C). Such strand bias suggests a possible relation of RepairSig.T2 with the transcription-coupled repair. In particular, a sub-pathway of the NER pathway, the so-called transcription-coupled NER (TC-NER) repairs the transcribed strand of transcriptionally active genes more efficiently than non-transcribed strand and transcriptionally silent DNA. However, additional experimental evidence is needed to support this hypothesis and, at this point, other mechanisms can not be excluded.

Finally, signature RepairSig.R2 is most similar to COSMIC Signatures 2 and 13 (Fig. 2B, cosine similarity: 0.71 and 0.67, respectively), both signatures known to be related to the mutagenic activity of the APOBEC family of cytidine deaminases. These enzymes convert cytosine to uracil in RNA and single-stranded DNA (ssDNA). Although both APOBEC signatures were included as the primary signatures in this analysis, RepairSig still recovered a secondary signature RepairSig.R2 as a composition of these two APOBEC signatures. This implies that the inferred signature has unique properties that cannot simply be modeled as a linear combination of Signatures 2 and 13. To understand the source of this non-additivity, we analyzed the local regional activity of RepairSig.R2 and Signatures 2, 13, and 3 with respect to replication timing. While the activities of Signatures 2 and 13 increase with replication time, the activity of RepairSig.R2 decreases (Fig. 2D,E). This suggests that early replicating regions provide a unique mutation opportunity for RepairSig.R2 that was not captured by Signatures 2 and 13. Since the APOBEC enzyme can only act on ssDNA, this opportunity has to be related to the cancer-specific emergence of ssDNA in early replicating regions. Our earlier studies revealed that APOBEC mutations in densely mutated, early replication enriched regions are associated with homologous recombination deficiency (HRD) [30,16]. Consistent with these observations, we hypothesize that RepairSig.R2 found in breast cancer is a result of ssDNA regions emerging in connection to HRD [20]. This interpretation is additionally supported by the fact that COSMIC Signature 3 associated with HRD has a similar local activity pattern to RepairSig.R2 (Fig. 2E).

### 2.3 Cross-cancer analysis of MMRd signatures

Next, we asked if the MMRd signature identified by RepairSig in BRCA can be observed in other cancer types as well. To this end, we applied RepairSig to additional cancers known to be affected by MMR deficiency: esophageal adenocarcinoma (ESAD), colorectal cancer (COCA), gastric cancer (GACA), and uterine corpus endometrial carcinoma (UCEC) (see Materials and Methods). Here, we use the model that is based on local genomic regions defined by the strand-specific direction of transcription. For GACA, ESAD and UCEC, we recovered a signature strikingly similar to the signature uncovered for BRCA (cosine similarity: 0.88, 0.87, and 0.67, respectively), which suggests it being a universal MMRd signature present in multiple cancer types.

To elucidate the remaining secondary signatures, we clustered them according to their RePrints [29] (Fig. 3). RePrint computes, for every trinucleotide, the conditional probabilities of three possible mutations of the central nucleotide under the assumption that this central nucleotide is mutated; it finds similarities between signatures independently of their mutational opportunities. These probabilities are represented as a vector called RePrint. RePrint similarity can point to signatures potentially sharing the same DNA repair deficiency even when these signatures share low cosine similarity [29] (Fig. 3). With exception of COCA, one of the secondary signatures uncovered in each cancer is likely to be MMRd-related, based on either a high cosine similarity with the reference *Δ*MSH6 or the RePrint similarity (Fig. 3). COCA displays two unique but distinct signatures. While the first is related to the HRD cluster and consistent with occurrences of HRD in colorectal cancer, the second does not reassemble any other signature. This might indicate a unique DNA repair deficiency but might also result from a missing primary signature from the model.

**Fig. 3.**
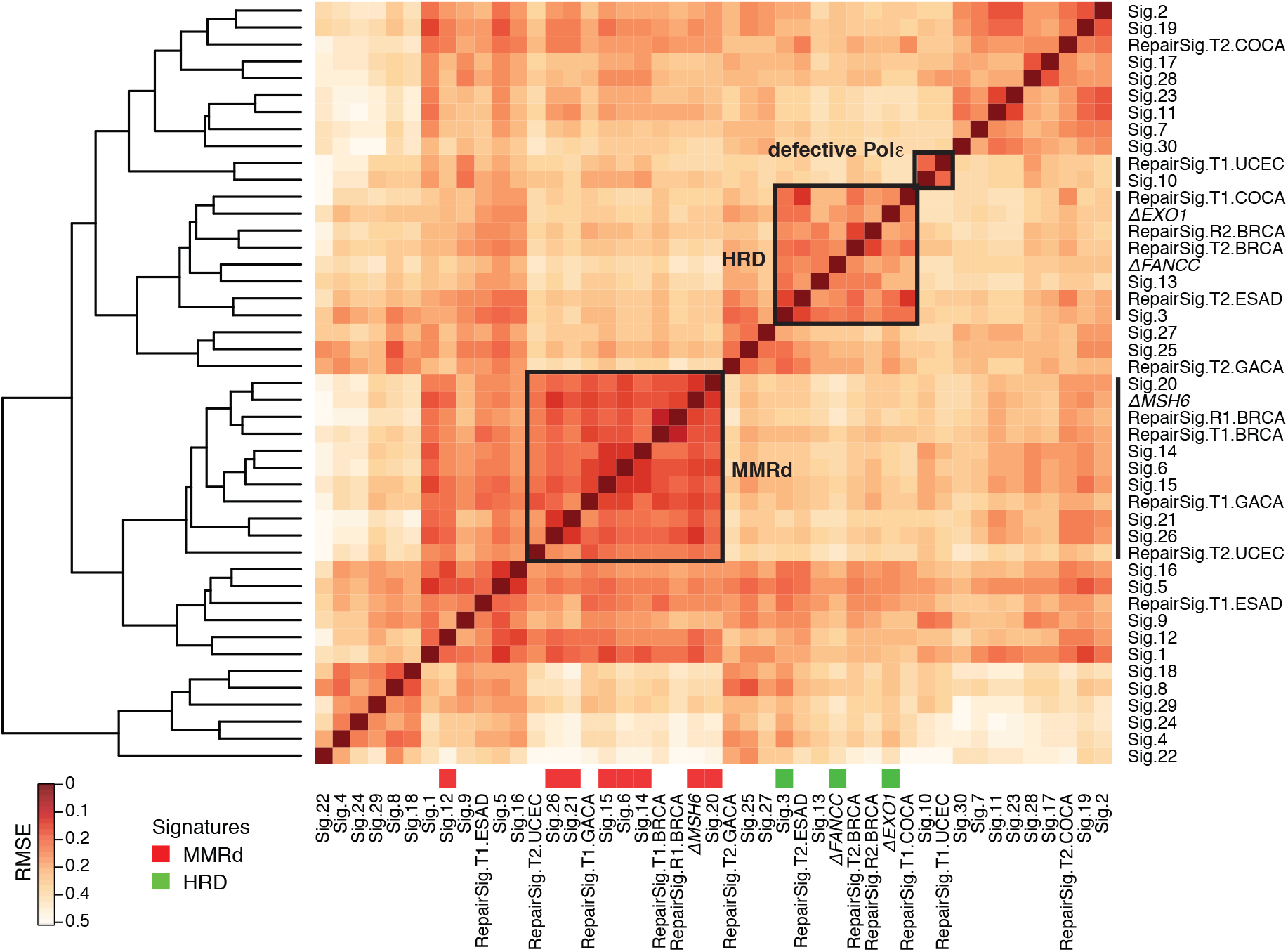
Similarities of inferred signatures. Clustering of RePrint of COSMIC mutational signatures, three signatures from knockout experiments of DNA repair genes: MSH6, FANCC, and EXO1, and signatures inferred by RepairSig across five cancer types. The signatures known to be related to MMR deficiency are marked by red squares below the heat map. Green squares mark signatures associated with HRD. Black boxes depict groups of highly similar RePrints of signatures related to the same process, as labeled. RMSE scale legend for signature RePrints is shown on the left.

## 3 Conclusions

In this study, we developed RepairSig, the first approach to infer DNA repair deficiency signatures that builds on a biologically realistic, non-additive model allowing for the composition of mutagenic processes. Because we model the mutational landscape as a composition of DNA damage and repair, only a single MMR signature was required to explain BRCA mutation data as opposed to several MMR signatures inferred by NMF-type models. In addition, the MMR signature inferred by RepairSig is more similar to the benchmark experimental MMR signature compared to any other COSMIC signatures and could be independently identified in several other cancer types.

Besides DNA repair deficiency, RepairSig can point to other non-additive contributions to the cancer landscape such as cancer-specific changes in mutation opportunity. In this paper, we focused on the case where signatures of primary processes are assumed to be known while signatures of secondary processes are to be inferred. However, RepairSig can be used to identify unknown primary processes as well.

It should be noted that in addition to pairwise interactions between primary and secondary mutagenic processes, more complex interactions are also possible. However, we limit our model to pairwise interactions only since inferring higher-order interactions would require significantly larger data sets. Nevertheless, our study provides a proof-of-principle for a new and powerful approach for discovering, analyzing, and interpreting mutational signatures in cancer.

## 4 Materials and Methods

### 4.1 Data

#### Cancer data

In this study, we analyzed single base substitutions from several datasets of cancer sequences that are known to be affected by the deficiency in DNA repair mechanism. The main dataset used in this study is the cohort of 560 breast cancer (BRCA) whole-genomes previously analyzed by Nik-Zainal et al. [23]. We also analyzed single base substitutions from the International Cancer Genome Consortium (ICGC Release 25) [3] in 30 colorectal (COCA), 265 esophageal adenocarcinoma (ESAD), and 31 gastric (GACA) whole-cancer-genome sequences. In addition, we used a subset of the endometrial carcinoma (UCEC) cohort downloaded from ICGC (Release 28); we selected 65 highly mutated whole-exome sequences from patients with mutations in POLE/POLD proofreading genes.

#### Genomic regions

To analyze mutations in genomic regions related to transcriptional strands, we downloaded the GRCh37 human gene annotations from the Ensembl database [2]. We assigned each mutation to the intergenic region, the transcribed (non-coding, template), or the non-transcribed (coding, non-template) strand based on the gene orientation and the strand-specific location of the pyrimidine base of the mutation. For replication time analysis in BRCA, we downloaded percentage normalized replication time estimates based on Repli-seq data in the MCF-7 cell line from the ENCODE project [1]. We split the genomic intervals into deciles based on Repli-seq estimates. To build the three-dimensional input matrix *M*, we counted the number of mutations falling into each region/domain (transcriptional or replicative) for each sample and mutation type separately. All results were corrected for the size of each genomic domain.

#### Mutational signatures

In this study, we used a set of 30 validated mutational signatures downloaded from the Catalogue of Somatic Mutations in Cancer (COSMIC, version 2) consortium [10] and three mutational signatures derived from cell line screens with targeted CRISPR-Cas9-based knockouts of DNA repair genes: MSH6, FANCC, and EXO1, from Zou et al. [31]. The primary signature matrix *P* consists of mutational signatures assumed being “active” in a given cancer type with the exception of signatures related to defective DNA repair processes (e.g. MMR deficiency signatures 6, 12, 14, 15, 20, 21, and 26) that are inferred in RepairSig as secondary signatures (matrix *Q*). In addition, we assume that *P* contains a *background* mutational process that represents spontaneous mutations that usually arise during DNA synthesis. Here, the background process was computed from the trinucleotide frequencies across the genome corresponding to mutation opportunities. However, the background process can also be inferred directly in our model.

To determine which signatures are active in a given cancer type, we either relied on published reports (BRCA [23], UCEC [5]) or used the quadratic programming approach with bootstrapping available in the R package – `SignatureEstimation` [14]. To detect signatures present in COCA, ESAD, and GACA cancers, we performed two rounds of signature filtering with `SignatureEstimation`. We started with the full set of 30 COSMIC mutational signatures, performed the bootstrapping procedure, removed any signatures that did not have sufficient statistical support (at least 1 sample with 0.1 contribution or 25% of samples with 0.05 contribution at a p-value threshold of 0.01), and repeated the bootstrapping procedure. The signatures that passed the second round were used in our analysis of COCA, ESAD, and GACA cancers.

### 4.2 Additional details of RepairSig

#### Tensor algebra formulation

The current approaches for modeling mutational processes are mostly focused on decomposing a catalog of mutations from cancer genomes (*M*) into a set of mutational signatures (*P*) and their sample-specific activities (*E*). It is usually achieved with NMF methods by solving a factorization problem *M* ≈ *A* × *P* (here only a single genomic region is considered). This problem is generalized by our RepairSig model to include local genomic properties of mutational processes as well as the joint effect of the primary and secondary mutational processes as explained in Section 2.1. To solve the new problem we took advantage of the tensor formalism and the powerful TensorFlow framework built for high-performance numerical computation and machine learning. The matrices defined in Section 2.1 can be expressed as third-order tensors, either directly in the case of the matrix **M**^*G*×*K*×*L*^ ≔ *M*^*G*×*K*×*L*^, or by introducing an extra (dummy) dimension **P**^*N*×*K*×1^ ≔ *P*^*N*×*K*^, where superscripts represent the dimensions and the ‘1’ represents the dummy dimension introduced for dimension compatibility. Here, we use bold and italic letters to distinguish tensors and matrices, respectively. We can now express Eq. (2) in tensor form as follows

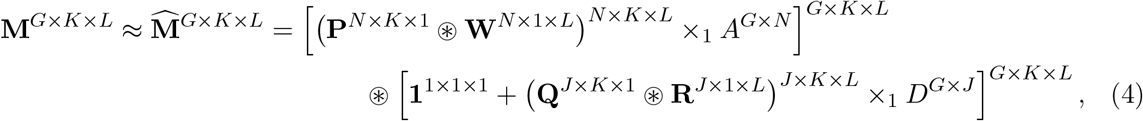

where ⊛ is the Hadamard (element-wise) product of two tensors and ×_1_ is the *n*-mode (*n* = 1) product of a tensor with a matrix (see review by Kolda & Bader [18]). For the Hadamard product, we make use of the NumPy style *broadcasting operation*, which is the process of making tensors with different shapes compatible for arithmetic operations. It broadcasts (copies) a tensor across a particular dimension so that the tensors have compatible shapes. The dimensions of all tensors, matrices, and tensor products are provided for clarity.

The goal is to factorize the input matrix *M* into lower dimension tensors and matrices as detailed by Eq. (4). We assume that the mutational signature matrix *P* for the primary mutational processes is known (predefined). Unknown and to be identified are the secondary mutational signature matrix *Q*, the exposure matrices *A* and *D* for the respective primary and secondary mutational processes, and the matrices *W* and *R* representing the local regional strength of mutational process activities. All tensors and matrices are non-negative and the sum of values for each signature in *P*, *Q*, *W* and *R* along *K* and *L* dimension normalizes to 1 (i.e. they are probability distributions). The acceptable solution should minimize the Frobenius norm between the input matrix *M* and its reconstructed approximation according to Eq. (4) and it should satisfy the specified constraints. Formally, we optimize the following optimization problem:

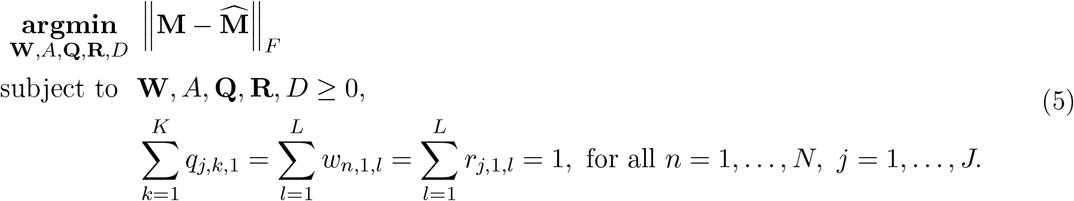

#### Implementation

Our method is implemented as an iterative gradient descent optimization problem using the Keras API in TensorFlow v2.x [4] using Python 3. As the loss function, we adopted the Frobenius norm between *M* and the right side of eq. (4) to evaluate the fit of our model to the given data. We chose the adaptive moment estimation method (Adam) [17] in combination with a custom learning rate strategy as our optimizer. Adam is well suited for problems with sparse data and achieved a suitable trade-off between performance and accuracy during benchmarks using the simulated data.

#### Model training

Starting with random tensors for *W*, *A*, *Q*, *R*, and *D*, in each iteration, the gradients for these tensors are computed based on the loss function. Consequently, these gradients are scaled according to the Adam optimizer using a user-specified learning rate and applied to the corresponding tensors. To reduce the probability of prematurely converging towards local minima during the gradient descent optimization, we bracket our optimization into a series of steps defined by learning rate and number of iterations. This allows to optimize our problem formulation by starting with an initially large learning rate and, as the optimization begins to converge, reduce the learning rate in consecutive steps. The particular choice for the learning rate and the number of iterations is dependent on the input data and may require manual adjustment.

Technically, Eq. (5) corresponds to a constraint optimization problem. To account for these constraints in TensorFlow, we apply an additional projection function after each optimization step designed to clip negative values to 0 hence satisfying the non-negativity constraints, and to normalize values for each signature in tensors *W*, *Q*, and *R*, so they sum to one, therefore, conforming to the probability distribution constraints. This, combined with the above-described optimization strategy enables RepairSig to leverage the optimization power offered by TensorFlow while still ensuring convergence towards a valid and biologically meaningful solution. All model parameters are learned from the input data in an unsupervised manner. All models derived in this study were trained on Nvidia V100 tensor core GPUs with 640 tensor cores and 32GB of memory.

## Acknowledgements

This study was supported by the Intramural Research Programs of the National Library of Medicine (NLM), National Institutes of Health, USA. This work utilized the computational resources of the NIH HPC Biowulf cluster (http://hpc.nih.gov). We thank Ariella Saslafsky for helpful discussions, language editing, and proofreading.

## Supplementary Materials

### S1 Statistical analysis

All statistical analyses and figures were done in R version 3.5.1. The heat maps of similarity between signature RePrints (Fig. 3) were generated with unsupervised hierarchical clustering of the matrix of root-mean-square error (RMSE) values computed between each pair of RePrints. We used the heatmap.2 function from R with default parameters (Euclidean distance and complete linkage method).

### S2 Simulated data

Generation of simulated data was required to rigorously test the performance of RepairSig with established ground truth signatures, their exposures, and local regional activities for both primary and secondary processes. The mutation count matrix *M* that serves as an input to our method was generated as specified by Eq. (4). In order to compute *M*, the six matrices *P*, *Q*, *A*, *W*, *R* and *D* used in Eq. (4) needed to be simulated as described below. These matrices were later used to test the performance of RepairSig. For this simulation, we assumed that *G* = 7000 (number of samples), *K* = 96 (number of mutation types), *L* = 10 (number of local genomic regions), *N* = 17 (number of primary mutational signatures), and *J* = 1,…, 3 (number of secondary mutational signatures).

#### Signature matrices *P* and *Q*

The signature matrix *P* for the primary mutational processes is composed of a total of *N* = 17 signatures consisting of 16 COSMIC mutational signatures (version 2) and the background signature (see the ‘Mutational signatures’ in Materials and Methods). The MMR deficiency signatures and signatures exhibiting a linear dependency with other signatures (Signatures 4, 5, 8, 9, 23, 25, and 29, see [14]) were excluded. The experimental signature *Δ*MSH6 [31] and COSMIC Signatures 6 and 12 were used to construct the signature matrix *Q* for the secondary mutational processes for *J* = 1,…, 3.

#### Exposure matrices *A* and *D*

To underscore the fact that in most cancers only a small amount of signatures are active in a single sample, we initialized all elements of matrices *A* and *D* to zero. For each signature, we then selected a small number of samples (between 20-30%), in which a given signature would be active, at random. Values for these activities (exposures) were drawn from a truncated normal distribution in the range of [100,10000] with mean *μ* = 4000 and variance *σ*^2^ = 2000. This procedure was performed separately for *A* and *D*.

#### Regional activity matrices *W* and *R*

Regional activity matrices *W* and *R* were initialized with random values drawn from a truncated normal distribution 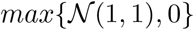. Then, for each signature, we selected a small number of regions with substantially increased and decreased activities (20-40% of regions for each type). For each signature *n* and each region *l*, we updated the local activity value *w_n,l_* according to the following equation:

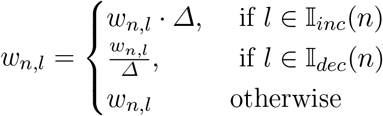

where 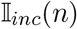 and 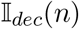 denote sets of genomic region indexes selected for increased and decreased activities (respectively) for signature *n*. New random value *Δ* was drawn from 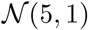 for each update of *w_n,l_*. Finally, the regional activities were normalized to sum up to 1 for each signature. Matrix *R* of local regional activities of the secondary mutational processes was generated analogously.

Summary of the performance of RepairSig on the simulated data.

**Fig. S1.**
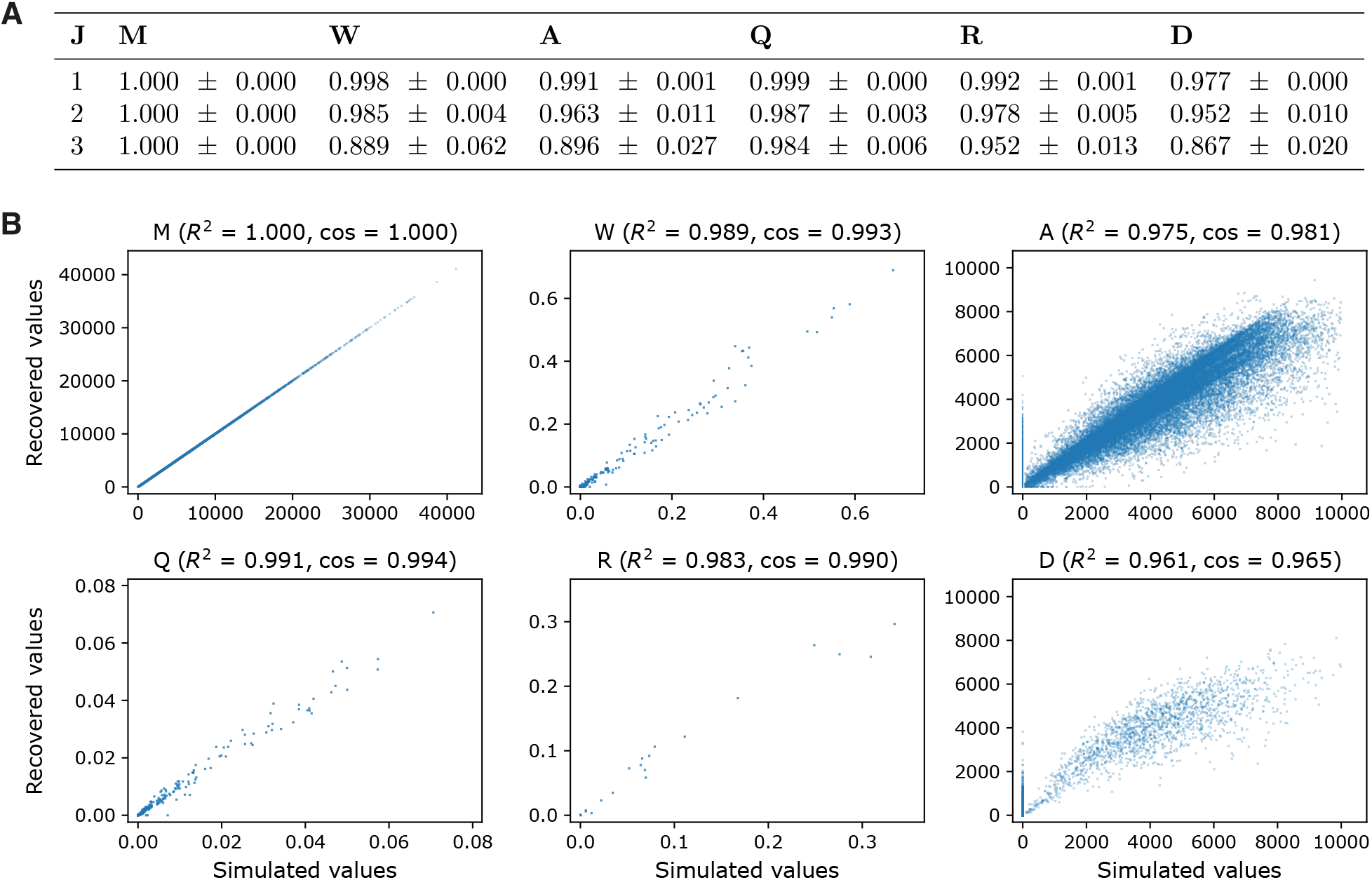
Evaluation of RepairSig on simulated data. **(A)** Pearson correlation, averaged over 10 optimizations, between the simulated data and RepairSigs solution for matrices *M*, *W*, *A*, *Q*, *R*, and *D* as well as three DNA repair signature sizes (*J* = 1, 2, 3). The standard deviation between the runs is provided in the second term. **(B)** Scatter plots between the simulated dataset and RepairSigs best solution with the lowest loss over 10 runs. Each data point corresponds to pair a (*x, y*) with values originating from elements at matching positions in the simulated matrix (x-axis) and the matrix recovered by our method (y-axis). Pearson correlation and cosine similarity are indicated in brackets for each matrix. Overall, RepairSig was able to recover all matrices with high accuracy.

**Fig. S2.**
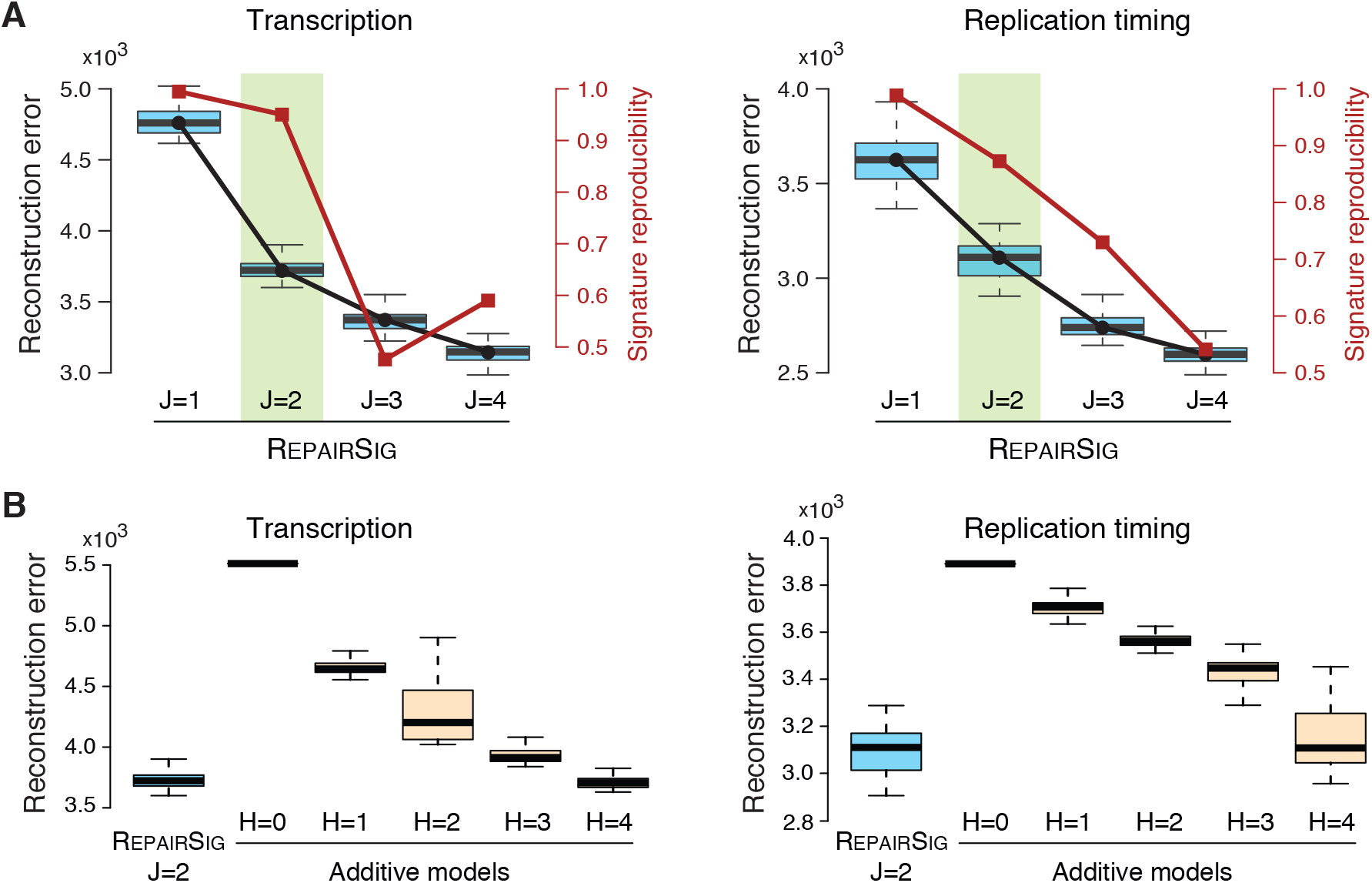
Model comparison on BRCA dataset. (**A**) Comparative assessment of RepairSig’s performance on the reconstruction error (black) and the signature reproducibility (red) for an increasing number of the secondary signatures (*J* = 1,…, 4). The reconstruction error was measured by the Frobenius norm of the difference between the input mutation count matrix *M* and its reconstructed approximation. The signature reproducibility was computed using the Silhouette score of the signature clustering into *J* groups from 100 random initialization runs of RepairSig using cosine similarity. For *J* = 1, we used the average cosine similarity between all signatures as a measure of the signature reproducibility. RepairSig with *J* = 2 (green) shows the best performance by balancing the reconstruction error and the signature reproducibility for both variants of RepairSig modeling local signature strength in the transcriptional and replication timing domains. (**B**) Comparison of the reconstruction error between the best RepairSig model (*J* = 2) and the classical model that assumes that all mutational signatures are additive (i.e. there are no secondary processes). For the additive model, we inferred *H* = 0,…, 4 mutational signatures that complemented the set of signatures being active in BRCA. For *H* > 0, all MMRd signatures active in BRCA have been removed from the set of predefined signature as they were being modeled by the inferred signatures. For *H* = 0, all 12 known signatures being active in BRCA have been used in the additive model and no signature was inferred.

## References

1. ENCODE: Encyclopedia of DNA Elements. https://www.encodeproject.org/, Accessed: 2020-01-31

2. Ensembl Genomes project. https://www.ensembl.org, Release: 99

3. International Cancer Genome Consortium (ICGC). https://dcc.icgc.org/

4. Abadi, M., Agarwal, A., Barham, P., Brevdo, E., Chen, Z., Citro, C., Corrado, G.S., Davis, A., Dean, J., Devin, M., Ghemawat, S., Goodfellow, I., Harp, A., Irving, G., Isard, M., Jia, Y., Jozefowicz, R., Kaiser, L., Kudlur, M., Levenberg, J., Mané, D., Monga, R., Moore, S., Murray, D., Olah, C., Schuster, M., Shlens, J., Steiner, B., Sutskever, I., Talwar, K., Tucker, P., Vanhoucke, V., Vasudevan, V., Viégas, F., Vinyals, O., Warden, P., Wattenberg, M., Wicke, M., Yu, Y., Zheng, X.: TensorFlow: Large-scale machine learning on heterogeneous systems (2015), https://www.tensorflow.org/, software available from tensorflow.org

5. Alexandrov, L.B., Kim, J., Haradhvala, N.J., Huang, M.N., Tian Ng, A.W., Wu, Y., Boot, A., Covington, K.R., Gordenin, D.A., Bergstrom, E.N., Islam, S.M.A., Lopez-Bigas, N., Klimczak, L.J., McPherson, J.R., Morganella, S., Sabarinathan, R., Wheeler, D.A., Mustonen, V., Getz, G., Rozen, S.G., Stratton, M.R., Alexandrov, L.B., Bergstrom, E.N., Boot, A., Boutros, P., Chan, K., Covington, K.R., Fujimoto, A., Getz, G., Gordenin, D.A., Haradhvala, N.J., Huang, M.N., Islam, S.M.A., Kazanov, M., Kim, J., Klimczak, L.J., Lopez-Bigas, N., Lawrence, M., Martincorena, I., McPherson, J.R., Morganella, S., Mustonen, V., Nakagawa, H., Tian Ng, A.W., Polak, P., Prokopec, S., Roberts, S.A., Rozen, S.G., Sabarinathan, R., Saini, N., Shibata, T., Shiraishi, Y., Stratton, M.R., Teh, B.T., V?zquez-Garc?a, I., Wheeler, D.A., Wu, Y., Yousif, F., Yu, W.: The repertoire of mutational signatures in human cancer. Nature 578(7793), 94–101 (02 2020)

6. Alexandrov, L.B., Nik-Zainal, S., Wedge, D.C., Aparicio, S., Behjati, S., et al.: Signatures of mutational processes in human cancer. Nature 500(7463), 415–421 (2013). https://doi.org/10.1038/nature12477, http://dx.doi.org/10.1038/nature12477

7. Alexandrov, L.B., Nik-Zainal, S., Wedge, D.C., Campbell, P.J., Stratton, M.R.: Deciphering Signatures of Mutational Processes Operative in Human Cancer. Cell Reports 3(1), 246–259 (2013). https://doi.org/10.1016/j.celrep.2012.12.008, http://dx.doi.org/10.1016/j.celrep.2012.12.008

8. Drost, J., Boxtel, R.v., Blokzijl, F., Mizutani, T., Sasaki, N., et al.: Use of crispr-modified human stem cell organoids to study the origin of mutational signatures in cancer. Science p. eaao3130 (2017). https://doi.org/10.1126/science.aao3130, http://dx.doi.org/10.1126/science.aao3130

9. Fischer, A., Illingworth, C.J., Campbell, P.J., Mustonen, V.: EMu: probabilistic inference of mutational processes and their localization in the cancer genome. Genome Biology 14(4), 1–10 (2013). https://doi.org/10.1186/gb-2013-14-4-r39, http://dx.doi.org/10.1186/gb-2013-14-4-r39

10. Forbes, S.A., Beare, D., Boutselakis, H., Bamford, S., Bindal, N., et al.: Cosmic: somatic cancer genetics at high-resolution. Nucleic Acids Research 45(D1), D777–D783 (2017). https://doi.org/10.1093/nar/gkw1121, http://dx.doi.org/10.1093/nar/gkw1121

11. Gonzalez-Perez, A., Sabarinathan, R., Lopez-Bigas, N.: Local Determinants of the Mutational Landscape of the Human Genome. Cell 177(1), 101–114 (03 2019)

12. Haradhvala, N.J., Kim, J., Maruvka, Y.E., Polak, P., Rosebrock, D., Livitz, D., Hess, J.M., Leshchiner, I., Kamburov, A., Mouw, K.W., Lawrence, M.S., Getz, G.: Distinct mutational signatures characterize concurrent loss of polymerase proofreading and mismatch repair. Nat Commun 9(1), 1746 (05 2018)

13. Haradhvala, N., Polak, P., Stojanov, P., Covington, K., Shinbrot, E., et al.: Mutational strand asymmetries in cancer genomes reveal mechanisms of dna damage and repair. Cell 164(3), 538–549 (2016). https://doi.org/10.1016/j.cell.2015.12.050, http://dx.doi.org/10.1016/j.cell.2015.12.050

14. Huang, X., Wojtowicz, D., Przytycka, T.M.: Detecting presence of mutational signatures in cancer with confidence. Bioinformatics 34(2), 330–337 (2018). https://doi.org/10.1093/bioinformatics/btx604, http://dx.doi.org/10.1093/bioinformatics/btx604

15. Kim, J., Mouw, K.W., Polak, P., Braunstein, L.Z., Kamburov, A., et al.: Somatic ERCC2 mutations are associated with a distinct genomic signature in urothelial tumors. Nature Genetics 48(6), 600–606 (2016). https://doi.org/10.1038/ng.3557, http://dx.doi.org/10.1038/ng.3557

16. Kim, Y.A., Wojtowicz, D., Sarto Basso, R., Sason, I., Robinson, W., Hochbaum, D.S., Leiserson, M.D.M., Sharan, R., Vadin, F., Przytycka, T.M.: Network-based approaches elucidate differences within APOBEC and clock-like signatures in breast cancer. Genome Med 12(1), 52 (05 2020)

17. Kingma, D.P., Ba, J.: Adam: A method for stochastic optimization (2017)

18. Kolda, T.G., Bader, B.W.: Tensor decompositions and applications. SIAM Review 51(3), 455–500 (2009). https://doi.org/10.1137/07070111X, http://dx.doi.org/10.1137/07070111X

19. Li, S., Crawford, F.W., Gerstein, M.B.: Using sigLASSO to optimize cancer mutation signatures jointly with sampling likelihood. Nat Commun 11(1), 3575 (07 2020)

20. Li, X., Heyer, W.D.: Homologous recombination in DNA repair and DNA damage tolerance. Cell Res 18(1), 99–113 (Jan 2008)

21. Ma, J., Setton, J., Lee, N.Y., Riaz, N., Powell, S.N.: The therapeutic significance of mutational signatures from DNA repair deficiency in cancer. Nat Commun 9(1), 3292 (08 2018)

22. Moldovan, G.L., D’Andrea, A.D.: How the fanconi anemia pathway guards the genome. Annu Rev Genet 43, 223–249 (2009)

23. Nik-Zainal, S., Davies, H., Staaf, J., Ramakrishna, M., Glodzik, D., et al.: Landscape of somatic mutations in 560 breast cancer whole-genome sequences. Nature 534(7605), 47–54 (2016). https://doi.org/10.1038/nature17676, http://dx.doi.org/10.1038/nature17676

24. Riva, L., Pandiri, A.R., Li, Y.R., Droop, A., Hewinson, J., Quail, M.A., Iyer, V., Shepherd, R., Herbert, R.A., Campbell, P.J., Sills, R.C., Alexandrov, L.B., Balmain, A., Adams, D.J.: The mutational signature profile of known and suspected human carcinogens in mice. Nat Genet 52(11), 1189–1197 (Nov 2020)

25. Rosales, R.A., Drummond, R.D., Valieris, R., Dias-Neto, E., da Silva, I.T.: signeR: an empirical Bayesian approach to mutational signature discovery. Bioinformatics 33(1), 8–16 (2016). https://doi.org/10.1093/bioinformatics/btw572, http://dx.doi.org/10.1093/bioinformatics/btw572

26. Sertic, S., Pizzi, S., Cloney, R., Lehmann, A.R., Marini, F., Plevani, P., Muzi-Falconi, M.: Human exonuclease 1 connects nucleotide excision repair (NER) processing with checkpoint activation in response to UV irradiation. Proc Natl Acad Sci U S A 108(33), 13647–13652 (Aug 2011)

27. Shiraishi, Y., Tremmel, G., Miyano, S., Stephens, M.: A Simple Model-Based Approach to Inferring and Visualizing Cancer Mutation Signatures. PLOS Genetics 11(12), e1005657 (2015). https://doi.org/10.1371/journal.pgen.1005657, http://dx.doi.org/10.1371/journal.pgen.1005657

28. Volkova, N.V., Meier, B., Gonz?lez-Huici, V., Bertolini, S., Gonzalez, S., V?hringer, H., Abascal, F., Martincorena, I., Campbell, P.J., Gartner, A., Gerstung, M.: Mutational signatures are jointly shaped by DNA damage and repair. Nat Commun 11(1), 2169 (05 2020)

29. Wojtowicz, D., Leiserson, M.D.M., Sharan, R., Przytycka, T.M.: DNA Repair Footprint Uncovers Contribution of DNA Repair Mechanism to Mutational Signatures. Pac Symp Biocomput 25, 262–273 (2020)

30. Wojtowicz, D., Sason, I., Huang, X., Kim, Y.A., Leiserson, M.D.M., Przytycka, T.M., Sharan, R.: Hidden Markov models lead to higher resolution maps of mutation signature activity in cancer. Genome Medicine 11, 49 (2019). https://doi.org/10.1186/s13073-019-0659-1, https://doi.org/10.1186/s13073-019-0659-1

31. Zou, X., Owusu, M., Harris, R., Jackson, S.P., Loizou, J.I., Nik-Zainal, S.: Validating the concept of mutational signatures with isogenic cell models. Nature Communications 9(1), 1744 (2018). https://doi.org/10.1038/s41467-018-04052-8, http://dx.doi.org/10.1038/s41467-018-04052-8

